# Human behavior and comfort during load carrying to autonomous mobile robot

**DOI:** 10.1101/2023.07.03.547589

**Authors:** Hideki Tamura, Taiki Konno, Shigeki Nakauchi, Tetsuto Minami

## Abstract

Interactions between humans and autonomous mobile robots (AMRs) are expected to grow in smart cities to improve logistics operations, such as depositing packages on AMRs for pickup on the street. However, the way that humans walk and pass objects to an AMR when approaching each other remains largely unknown. We conducted two psychophysical experiments to clarify the behavior and comfort of humans when carrying a package and placing it on an AMR for load carrying. Participants were asked to approach a programmed AMR and pass the package in two experiments: 1) changing the stop distance and AMR speed and 2) changing the stop distance and package weight. Motion trackers quantified the participants’ walking speed and frequency of hesitation to walk. In addition, the subjective heaviness and comfort were recorded through a questionnaire during each trial. The results indicated that the participants’ speed decreased and hesitation probability increased when the stop distance of the AMR decreased. Nevertheless, the participants felt more comfortable with the close approach, whereas the package weight did not affect their behavior. By contrast, they felt uncomfortable when AMR remained still. These findings suggest that humans regard the AMR approach as load-carrying assistance and not as invading their personal space. To achieve a comfortable interaction in load carrying from humans to AMRs, we suggest that the AMR can closely approach a person without eliciting personal space invasion.

## 1 Introduction

We lift and lower objects daily to carry them to desired locations [1]. This task is at the core of many activities, and a recent study has suggested its importance from an ecological perspective [2]. Thus, lifting or carrying packages to reduce laborers’ burden has been studied in ergonomics, including the cases of carrying loads on the back [3–5] and by hand in front [6–8]. Additionally, various studies have been focused on carrying loads with agents such as robots in human–robot interactions [9]. For example, a human and a robot arm can pass any common object to each other [10–13]. Furthermore, a recent study reported that a deep neural network can estimate the physical load levels of workers using wearable inertial sensors [14].

The frequency of interactions between humans and autonomous mobile robots (AMRs) is expected to increase in daily living [15–17] because governments and companies strongly push forward the development of smart cities to, among other goals, address various concerns of logistics [18–21]. For example, on the street, people may receive packages from AMRs for last-mile delivery as well as deposit packages to AMRs for efficient pickup. Many studies on passing objects between humans and AMRs have been conducted [13, 22–24]. Although the robotics aspects have been mainly addressed, the human side of the interaction should also be studied to understand real operations in smart cities. In fact, we should analyze the human behavior and subjective responses, particularly regarding safety [25–28] and comfort [29]. For example, Lasota et al. [30] indicated that even when humans reach physical safety, they can feel frustration [31] or discomfort in certain situations, and simply preventing physical contact cannot guarantee a safe and comfortable interaction between humans and robots. Thus, the evaluation of human responses is required to develop useful AMRs and socially assistive robots [32–34].

As mentioned above, although various studies have demonstrated AMRs approaching and passing objects to humans, the human behavior when humans approach and pass objects to AMRs while both are moving remains to be understood. The human behavior when receiving an object is not necessarily the same as that when passing an object. Therefore, we focused on how humans pass an object to an AMR. First, we considered distance and speed, like in proxemics, because the distance between a human and a robot is essential for comfortable interactions [35, 36] and appropriateness [37]. For example, Torta et al. investigated the closeness of a person for comfortable communication with a small humanoid robot [38]. They reported a comfortable distance of 173 cm for standing participants, which was larger than the personal space (45–120 cm) for human-to-human communication [36, 39, 40]. Additionally, human–robot interactions have been explored regarding collision avoidance and focusing on walking trajectories or avoidance directions [28, 41–45]. However, the distance between a person and AMR and the speed required to achieve a comfortable interaction when a human passes an object to the AMR remain to be determined in realistic scenarios. Second, we included the package weight as an experimental condition. The weight when carrying loads may influence the distance and speed of approaching humans and AMRs as well as the perceived safety and comfort during human–robot interaction. Studies have been conducted on the extent to which humans perceive the weight of a lifted object regarding kinematic analysis [7] and comparisons between participants with and without chronic low back pain [46]. Regarding perception, humans expect an object to be lighter when lifted together than when lifted alone [47], and social power involves weight estimation [48]. Various studies have reported that the size of an object is perceived to affect its weight, as we generally feel large objects to be lighter than equally weighted small objects [49–51]. However, changes in weight perception when humans pass objects to AMRs are unclear as well as their effect on comfort.

In this study, we aimed to clarify the behavior and subjective comfort of humans when passing a load to an approaching AMR. In Experiment 1, we focused on the stop distance and speed of the AMR approaching participants and investigated how they influence human walking speed and behaviors. Specifically, the participants lifted, carried, and placed a package on an approaching AMR, which was programmed to stop at various distances. Motion trackers recorded the positions and velocities during the trials. In Experiment 2, we investigated the effects of the AMR stop distance and package weight on the perceived weight and comfort.

## 2 Experiment 1

In Experiment 1, we investigated the influence of the stop distance and the moving speed of the AMR on human behavior for a load-carrying task. We assembled an environment for a human psychophysical evaluation with AMRs and a motion measurement system of commercially available virtual-reality hardware.

### 2.1 Methods

#### 2.1.1 Participants

Twenty healthy students (male; age, 22.4 ± 1.3 years) with normal or corrected-to-normal vision from Toyohashi University of Technology participated in this experiment. The recruitment for the experiment did not specify any gender preference. The sample size was determined based on a previous study [7], in which 20 participants were enrolled for a manual material-handling experiment. A power analysis using PANGEA [52] with sample size *n* = 20, effect size *d* = 0.45, and 5 replicates (see Section 2.1.3) revealed a power of 0.88. Nineteen participants were right-handed, with an average hand preference of 9.1 ± 1.5, and one was left-handed with a hand preference of −10, as assessed using the Flinders Handedness survey [53, 54]. Not all participants knew how the AMR carried an object, and they were new to performing an object-carrying task interacting with an AMR (see Section 2.1.2). All the experimental protocols involving human participants were approved by the Institutional Review Board of Toyohashi University of Technology, and the experiments were conducted in accordance with the Declaration of Helsinki. Written informed consent was obtained from all participants for the publication of their details.

#### 2.1.2 Apparatus

Fig. 1A illustrates the setup for Experiment 1. We ran the experiment in a square field (5 m × 5 m) separated by partitions with a height of 230 mm inside a room. The experimental environment contained only one participant and one AMR at a time, and they were placed 4 m apart. A computer for AMR control and another one for the trackers were placed, and the packages to be carried were placed on a table near the side of the experimenter. The experimenter operated the control and tracking systems outside the environment.

**Fig. 1.**
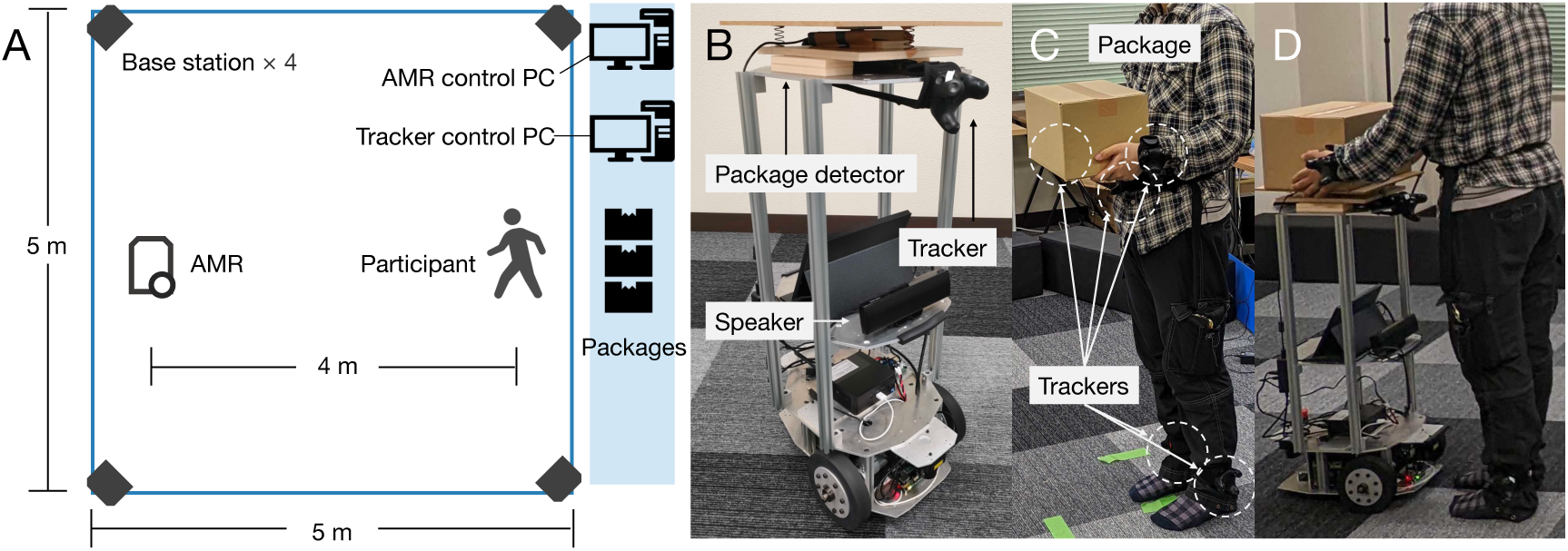
Setup for Experiment 1. **A** Environment. **B** AMR. **C** Motion trackers monitoring participant holding a package. **D** Participant passing package to AMR.

We used a custom-made wheeled platform (Mega Rover Ver 2.1, Vstone) as the AMR (Fig. 1B) with dimensions of 460 mm × 320 mm × 870 mm (length × width × height). The AMR used an onboard computer to control two motors directly linked to its wheel axles. A sensor on top of the AMR detected whether an object was placed. An additional computer was installed in the experimental environment to receive signals from the tracking system. These signals were then transmitted to the AMR onboard computer using a local Wi-Fi connection. The Robot Operating System (Melodic) running in Ubuntu 18.04 LTS controlled both computers.

Each participant wore five motion trackers (VIVE Tracker 3.0, HTC) on the waist, wrists, and ankles to measure motion (Fig. 1C). The AMR was equipped with a tracker on top of the front side. A computer tracking system running Windows 10 established communication with the trackers under the control of Unity (2020.3.20f1) and Steam VR (1.20.4), which are a development environment and software platform for virtual reality, respectively. Four base stations (Steam VR Base Station 2.0) in the experiment room defined the play area, and the system measured the six-axis position and orientation of each tracker at a 90 Hz sampling rate.

Ten cardboard boxes (320 mm × 230 mm × 200 mm) containing water bottles (1.35 kg as measured using an SH-20KN digital scale, A&D) were placed on the table in the experiment room.

#### 2.1.3 Procedure and tasks

Before conducting the experiment, the experimenter gave instructions to the participant and obtained written informed consent. In the initial state of each trial, the participants held a package with their hands (Fig. 1C) and stood at a distance of 4 m from the AMR. First, a speaker installed on the AMR emitted a buzzing sound as a cue to initiate the trial. The participants then started walking at their own comfortable pace from the designated starting point toward the AMR. When the participant took a step, the AMR began to move in a straight path toward the participant at a constant speed (low speed, 0.25 m/s; high speed, 0.5 m/s) until reaching a stop distance between the participant and AMR of 1, 2, 3, or 4 m. The last condition (4 m) indicated that the AMR did not move. The participants were asked to approach the AMR and place the package on the AMR at a distance at which they could behave naturally (Fig. 1D). Subsequently, the participant returned to the starting point, and one trial was completed. After the trial, the participants took a new package from behind and transitioned to the initial state for the next trial.

Each trial was randomly selected from all the combinations of stop distances between the AMR and participant (1, 2, 3, and 4 m) and AMR approaching speed (0.25 and 0.5 m/s). Trials with identical conditions were repeated five times. Thus, each participant completed 40 trials (5 trials × 4 distances × 2 speeds) in random order. The participants could take breaks at any time outside a trial.

#### 2.1.4 Data analysis

The signals from the motion trackers were analyzed using MATLAB R2022b (MathWorks). We excluded 17 trials corresponding to 2.1% of all trials because 1) they had over 10 missing values, 2) the initial buzzer sounded before the participant was ready, or 3) the AMR could not detect the package placement.

We defined the initial point of the participant as zero and the positive direction as that pointing toward the AMR. To correct for differences between trial times, we normalized them over time in percentages, from the instant of buzzing (0%) to that of package detection (100%) by the AMR. The participant’s walking speed was computed by dividing the Euclidean distance between each sampling of the participant’s waist tracker by the period between samples. Subsequently, a 1 Hz lowpass filter was applied (*lowpass* function of MATLAB with a steepness of 0.99) to remove high-frequency speed changes in a gait cycle because our cadence is approximately 100 steps per minute [55].

Statistical analyses were performed using JASP version 0.16.2 [56] and R version 4.0.4. The *p*-values were adjusted using Holm correction for multiple comparisons [57]. We calculated the adjusted degrees of freedom using the Greenhouse–Geisser correction if Mauchly’s test failed.

### 2.2 Results

#### 2.2.1 Position and speed

We computed the positions and speeds of the participants based on the location of the waist-attached motion tracker. Figs. 2A and 2B show the changes in the participants’ positions under the low and high AMR speeds, respectively. The curves show inverted U-shapes obtained by the participant approaching the AMR and then moving away from it. The peak, that is, the position where the participant slowed down, depended on the stop distance to the AMR. In addition, the changes were greater under the high AMR speed. The participants’ average trial time was 9.47 ± 1.03 s.

**Fig. 2.**
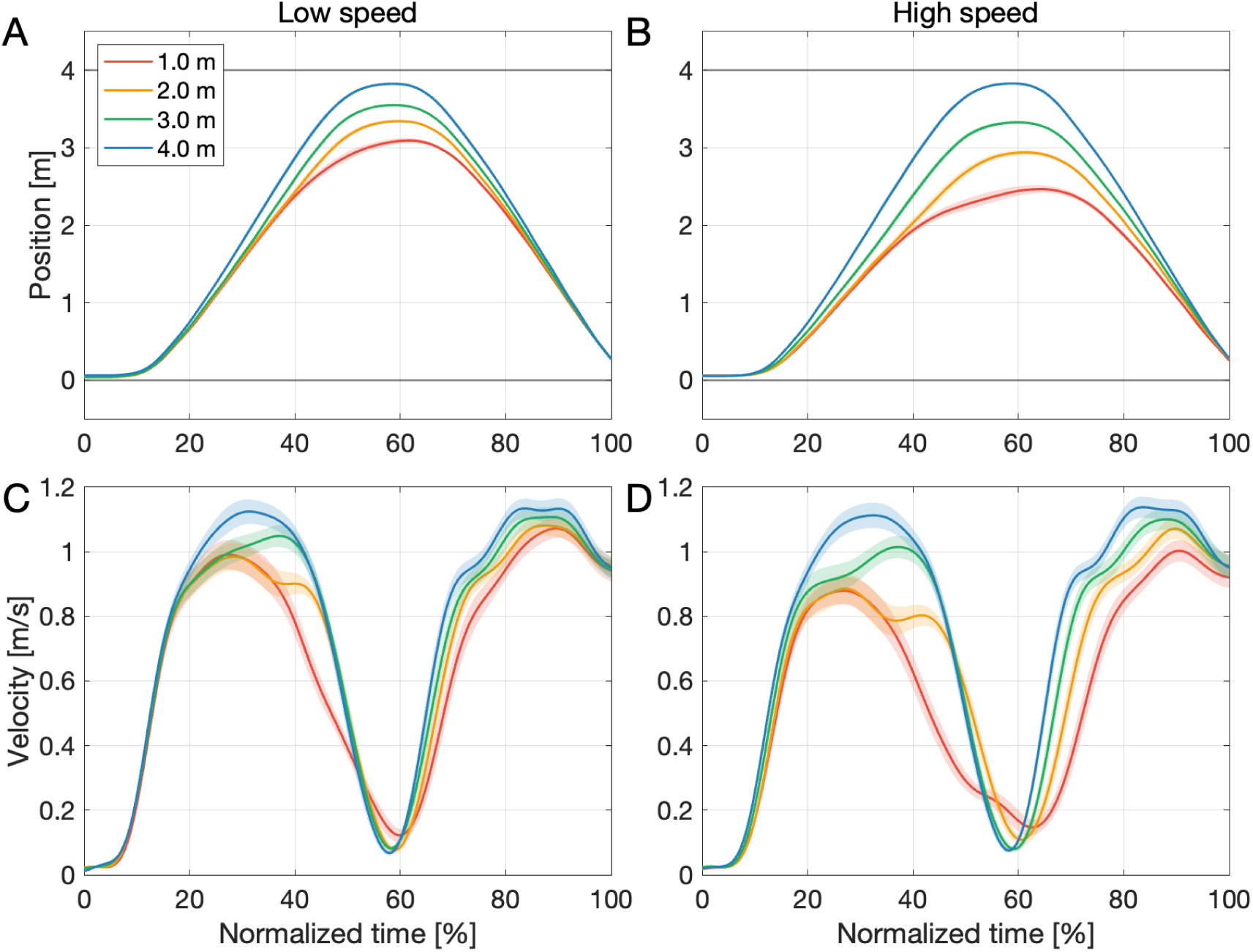
Positions and speeds of participants according to normalized time in Experiment 1. Positions under **A** low and **B** high AMR speeds. Speeds under **C** low and **D** high AMR speeds. The curve colors indicate the stop distances. The curves and shaded regions indicate the mean and standard errors across participants, respectively.

Figs. 2C and 2D show the changes in the participants’ walking speeds under the low and high AMR speeds, respectively. The walking speed increased to approximately 1.0 m/s and then decreased to approximately 0.1 m/s when the participant placed the package on the AMR. Then, the participant accelerated again when returning to the initial position.

Fig. 3 shows the maximum walking speed of the participants until the package was placed on the AMR as a quantitative indicator to compare the distance and speed. A two-way repeated- measures analysis of variance (ANOVA) indicated the main effects of distance (*F*(3, 57) = 44.7, *p* < 0.001, partial *η*^2^ = 0.702) and speed (*F*(1, 19) = 38.6, *p* < 0.001, partial *η*^2^ = 0.670). There was a significant interaction (*F*(3, 57) = 5.40, *p* < 0.005, partial *η*^2^ = 0.221). A post hoc test revealed that the speeds were higher in the following order: 1 m and 2 m < 3 m < 4 m (1 m vs. 2 m, *pholm* = 0.806; others, *pholm* < 0.001). Under the same distance conditions, the maximum walking speed under the high AMR speed was higher than that under the low AMR speed at 1 m and 2 m stop distances (*pholm* < 0.001 and *pholm* < 0.001, respectively) but not at 3 m and 4 m (*pholm* = 0.066 and *pholm* = 1.000).

**Fig. 3.**
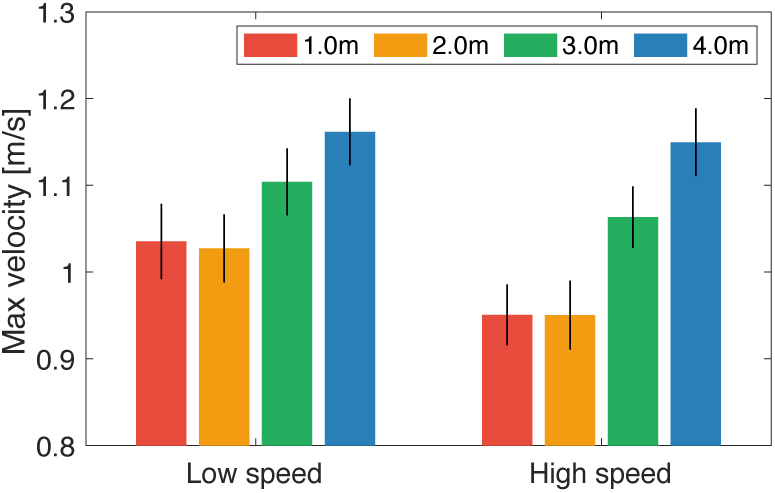
Average maximum speed of all participants in Experiment 1. The maximum speed was calculated between the trial initiation and package placement on the AMR. The bars indicate the standard errors.

#### 2.2.2 Hesitation for approaching

We found that participants sometimes hesitated to approach the AMR, particularly when the AMR speed was high and the distance from the participant was short. The participants slowly approached or temporarily stopped walking in these cases. These behaviors appeared as changes in the walking speed (solid line in Fig. 4A compared with dashed line). If we quantify these behaviors and compare their frequencies across the conditions, the human load-carrying behavior can be characterized. We defined a decision interval to determine whether hesitation occurred from 2 s after the trial began until 1 s before the package was placed on the AMR. During this interval, if the walking speed temporarily slowed, that is, if the local minimum was equal to or less than 50% of the maximum speed (i.e., sufficiently slow walking), we considered that the participant hesitated.

**Fig. 4.**
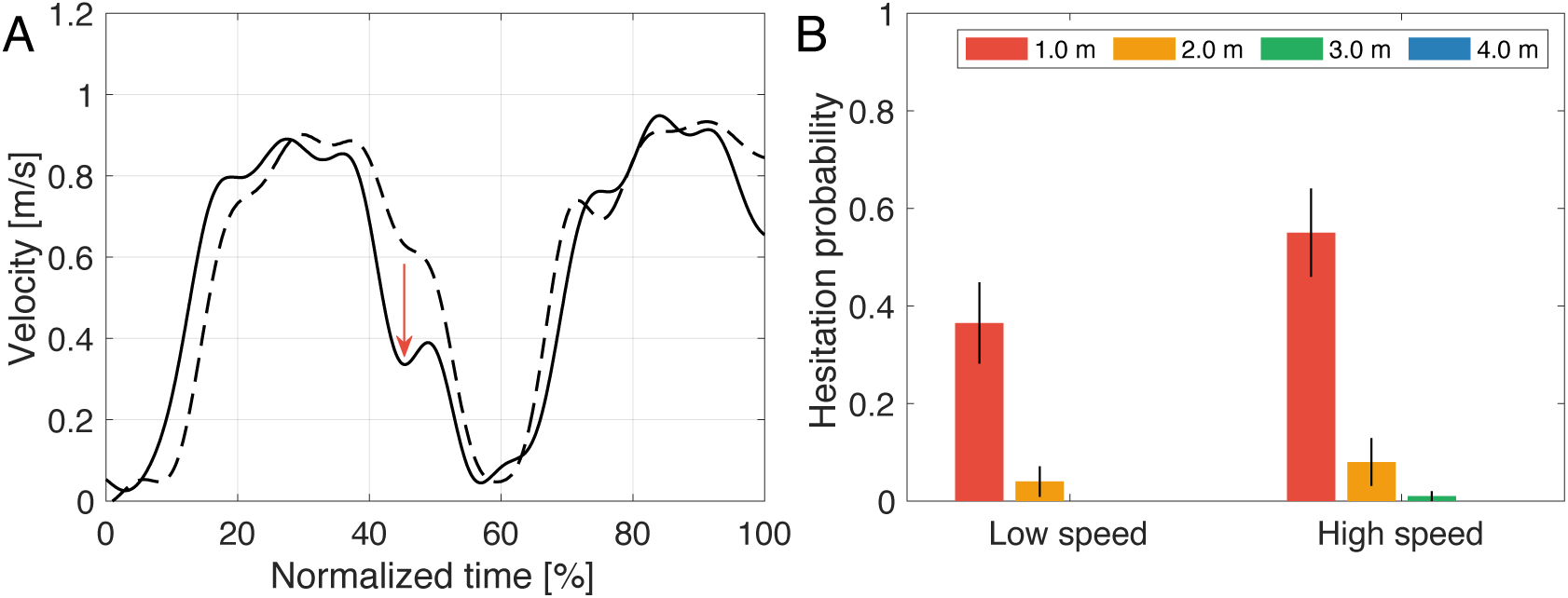
Hesitation probability in Experiment 1. **A** Sample trials showing hesitation (thick line with a red arrow indicating a point of rapid slowing) and its absence (dashed line) under the same condition (low AMR speed and 1 m stop distance) for different participants. **B** Average hesitation probability across participants under low and high AMR speeds.

We calculated the hesitation probabilities per condition and compared them, as shown in Fig. 4B. We performed a two-way repeated-measures ANOVA and observed the main effects of distance (*F*(1.38, 26.21) = 25.5, *p* < 0.001, partial *η*^2^ = 0.573) and speed (*F*(1, 19) = 12.9, *p* < 0.005, partial *η*^2^ = 0.405). A significant interaction (*F*(1.20, 22.81) = 5.82, *p* < 0.05, partial *η*^2^ = 0.234) was observed. A post hoc test indicated that the 1 m stop distance had a significantly higher hesitation probability than the other distances (1 m > 2 m, 3 m, 4 m; every *pholm* < 0.001), and the high AMR speed increased hesitation compared with the low speed (*pholm* < 0.005).

### 2.3 Discussion

Regarding position and speed, the position at which the participant placed the package on the AMR was closer to the initial point corresponding to the AMR stop distance and speed (Fig. 2). This suggests that the walking distance of the participants decreased when the AMR approached closer.

The maximum walking speed of the participants was higher when the stop distance was shorter (Fig. 3), and participants slowed when the AMR speed was higher. These results suggest that the pace-down caused participants’ discomfort with the AMR. For example, the distance eliciting an uncomfortable feeling is shorter when an object approaches a person than when the person approaches the object [58]. In Experiment 1, the AMR approached the participants (except at 4 m) and likely caused discomfort. We suggested that participants maintained a distance and slowed their pace to avoid feeling uncomfortable.

We also found that the hesitation probability was significantly higher at a 1 m stop distance (Fig. 4B). This suggests that participants hesitated when the AMR invaded their personal space. Humans feel more uncomfortable with robots operating at 1–3 m than at shorter or longer distances [59]. This range of uncomfortable distances is consistent with our findings, as indicated by the significantly lower maximum walking speed in Experiment 1. This phenomenon has been discussed in terms of personal space [36, 60]. Overall, the approaching AMR affected the personal space, walking speed, and hesitancy.

## 3 Experiment 2

The results of Experiment 1 showed that the participants slowed down and hesitated for shorter stop distances and higher AMR speed, suggesting that some discomfort with the AMR may occur. In Experiment 2, we further investigated whether the AMR operation affected comfort by letting the participants report their comfort level after the load-carrying task. In addition, we tested the influence of package weight on comfort. Based on the results of Experiment 1, we hypothesized that the participants’ comfort level would decrease when the AMR approached closer. Furthermore, as the walking speed of humans decreases when carrying a heavy load [61, 62], we hypothesized that the participants would walk slowly when carrying a heavier package, even in the presence of the AMR.

### 3.1 Methods

#### 3.1.1 Participants

Like in Experiment 1, we recruited twenty healthy students (18 males; age, 21.2 ± 1.8 years) from Toyohashi University of Technology who did not participate in Experiment 1 and had normal or corrected-to-normal vision. The power analysis using PANGEA [52] with *n* = 20, *d* = 0.45, and 3 replicates (see Section 3.1.3) revealed a power of 0.85. The participants were all right-handed with an average hand preference of 9.5 ± 1.1, as assessed using the Flinders Handedness survey [53, 54]. As for Experiment 1, we received approval from the Institutional Review Board of Toyohashi University of Technology, followed the Declaration of Helsinki, and obtained written informed consent from the participants.

#### 3.1.2 Apparatus

The environment, AMR control system, and tracking system were identical to those used in Experiment 1. We additionally placed 1) a monitor to indicate which package the participant carried and 2) a laptop computer to collect the survey responses behind the participant’s initial position. Moreover, the weights of the packages measured by the digital scale were 1.35 kg, 5.24 kg, and 10.24 kg, which we expected to be sufficiently distinguishable by the participants and remain within the weight range of common packages carried on a daily basis. The two heavier packages contained metal weights instead of water bottles. To mitigate the effect of size-weight illusion [49–51], the package appearances were identical, and the packages were labeled A, B, and C on their side faces for the participants to know the package to pick. There were four packages per weight, resulting in 12 packages in the environment. The participants did not know the weights of the packages.

#### 3.1.3 Procedure and task

First, the monitor displayed instructions to the participant, indicating which package they should pick from the table. Subsequently, the same load-carrying task as in Experiment 1 was performed. The stop distances were 1, 2, 3, and 4 m, while the AMR speed was fixed to 0.5 m/s, corresponding to the high speed in experiment 1. After each trial, the participants were asked the following two questions and provided their responses on a seven-point Likert scale through a spreadsheet shown on the laptop: Q1) How heavy did you feel the package was in this trial? (1, very light; 2, moderately light, 3, light; 4, neither light nor heavy; 5, heavy; 6, moderately heavy; 7, very heavy); Q2) How comfortable did you feel with the AMR operation in this trial? (1, very uncomfortable; 2, uncomfortable; 3, slightly uncomfortable; 4, neutral; 5, slightly comfortable; 6, comfortable; 7, very comfortable). After responding, the monitor instructed the next package to pick to begin a new trial. Each participant completed 36 trials (3 trials × 4 distances × 3 weights) in random order. The participants could take breaks at any time outside a trial.

#### 3.1.4 Data analysis

Preprocessing was performed as in Experiment 1. We omitted the data of one participant owing to a hardware issue and failure to collect the data and those of other three participants because they missed three trials in various conditions, impeding to perform two-way ANOVA. Thus, 136 trials, corresponding to 18.9% of all the trials, were excluded from the analysis. The maximum walking speed and hesitation probability were calculated as in Experiment 1. The subjective responses were averaged across participants.

### 3.2 Results

Fig. 5 shows the changes in position (Figs. 5A–5C) and speed (Figs. 5D–5F) according to package weight. Overall, the position and speed exhibited the same trends as in Experiment 1. The top of the reverse U-shaped curve was longer when the package was heavier. In addition, the speed decreased more rapidly for heavier packages. The average trial time was 10.64 ± 1.53 s.

**Fig. 5.**
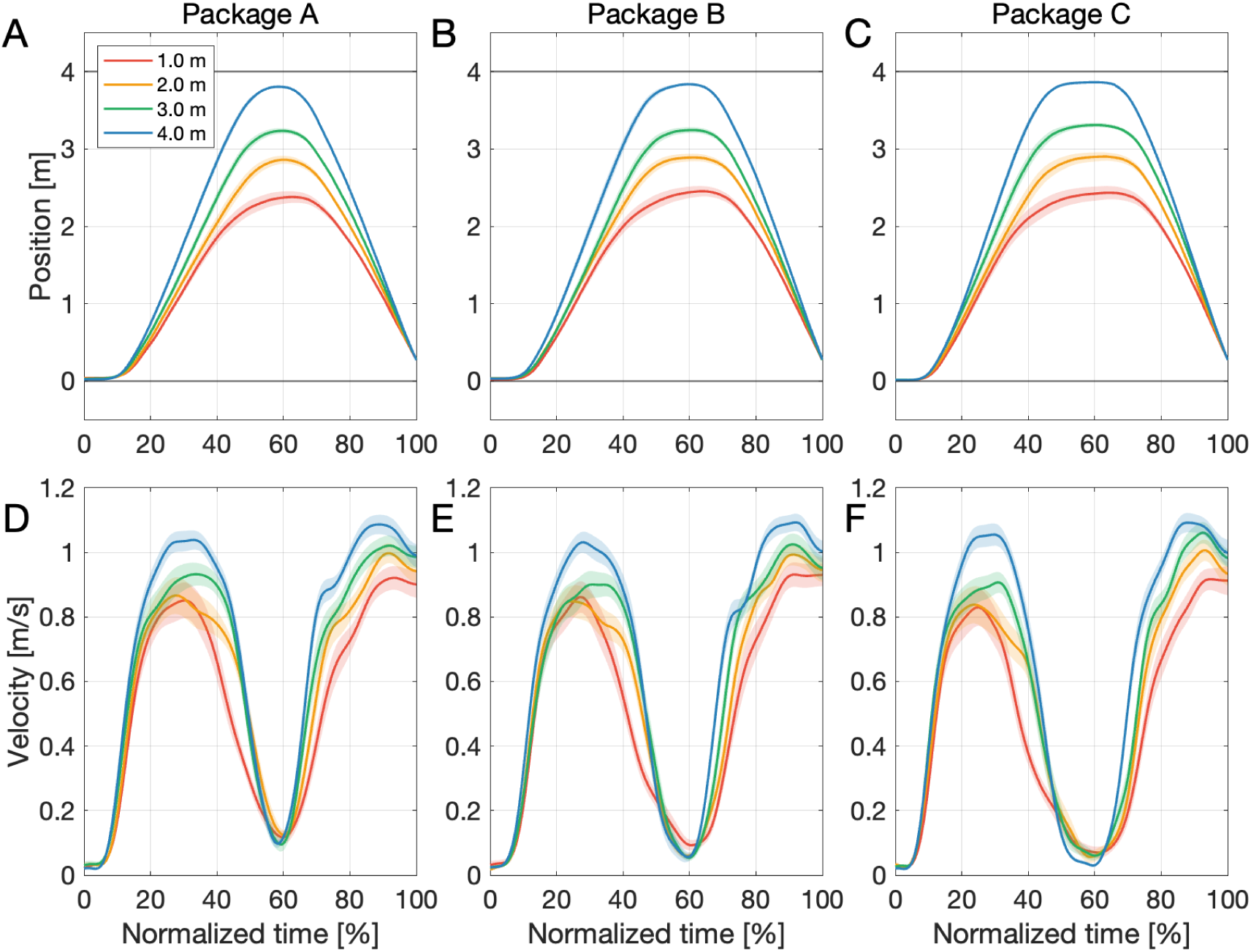
**A–C** Positions and **D–F** speeds of participants for different package weights (A: 1.35, B: 5.24, and C: 10.24 kg) in Experiment 2. The curve colors indicate the stop distances. The curves and shaded regions indicate the mean and standard errors across participants, respectively.

We compared the maximum speeds and hesitation probabilities between the different package weights, as shown in Figs. 6A and 6B. For the maximum speed, we performed a two-way repeated-measures ANOVA, and the main effect was significant for distance (*F*(1.22, 18.31) = 26.76, *p* < 0.001, partial *η*^2^ = 0.641). A post hoc test revealed that the maximum speed was lower when the stop distance was shorter (1 m < 2 m < 3 m < 4 m; 1 m vs. 2 m, *pholm* = 0.147; 2 m vs. 3 m, *pholm* = 0.146; 1 m vs. 3 m, *pholm* < 0.01; others, *pholm* < 0.001). By contrast, no main effect of weight (*F*(2, 30) = 1.97, *p* = 0.158, partial *η*^2^ = 0.116) or significant interaction (*F*(2.37, 35.60) = 0.83, *p* = 0.462, partial *η*^2^ = 0.052) was observed.

**Fig. 6.**
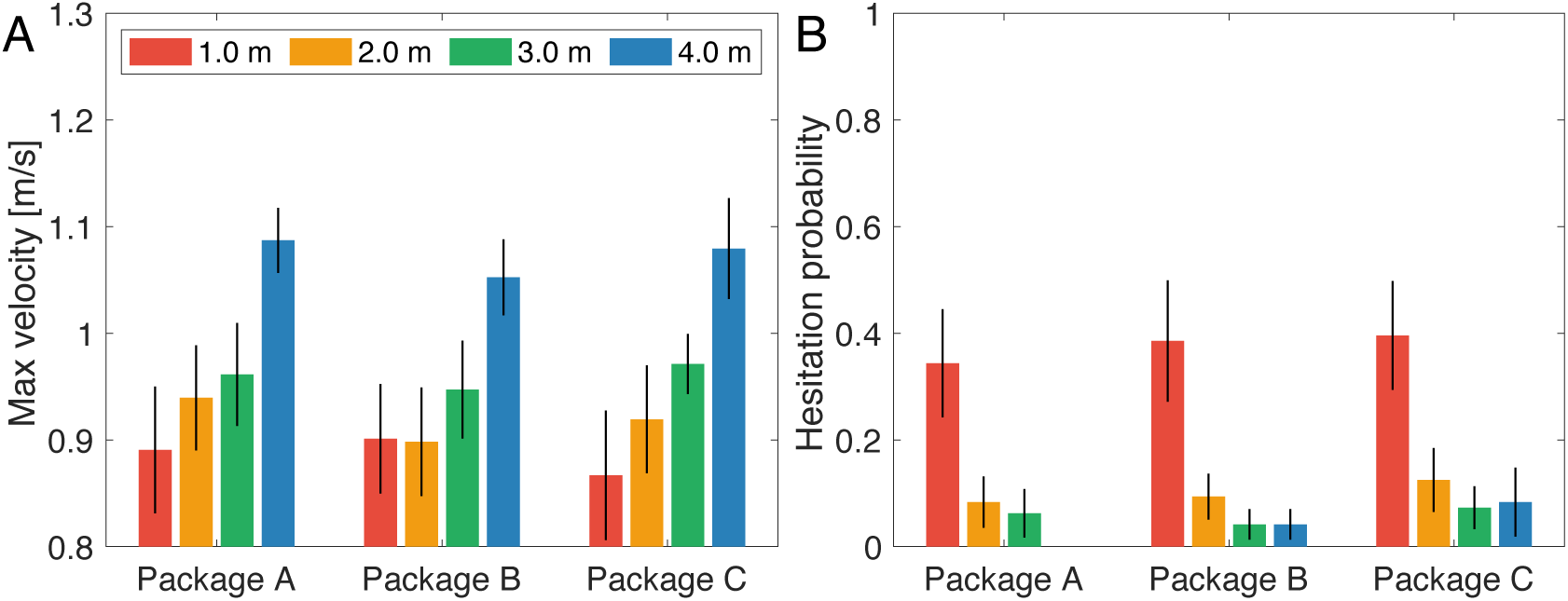
**A** Average maximum speed and **B** hesitation probability of all participants for different package weights in experiment 2. The bars indicate the standard errors.

Additionally, we performed a two-way repeated-measures ANOVA on the hesitation probabilities. There was a main effect of distance (*F*(1.41, 21.11) = 10.79, *p* < 0.005, partial *η*^2^ = 0.418), and the post hoc test revealed that a stop distance of 1 m had a significantly higher hesitation probability than the other distances (1 m < 2 m, 3 m, 4 m; every *pholm* < 0.001). No main effect of weight (*F*(1.45, 21.68) = 1.01, *p* = 0.356, partial *η*^2^ = 0.063) or significant interaction (*F*(3.25, 48.80) = 0.26, *p* = 0.870, partial *η*^2^ = 0.017) was observed.

Fig. 7A shows the average heaviness ratings obtained from the responses to question Q1. Two-way repeated-measures ANOVA showed a main effect of weight (*F*(2, 30) = 333.26, *p* < 0.001, partial η^2^ = 0.957). A post hoc test revealed that subjective heaviness was significantly higher when the packages were heavier (A < B < C; every *pholm* < 0.001), corresponding to the actual package weights. By contrast, no main effect of distance (*F*(1.56, 23.37) = 1.83, *p* = 0.188, partial *η*^2^ = 0.109) or significant interaction (*F*(4.05, 60.80) = 0.95, *p* = 0.442, partial *η*^2^ = 0.060) was observed.

**Fig. 7.**
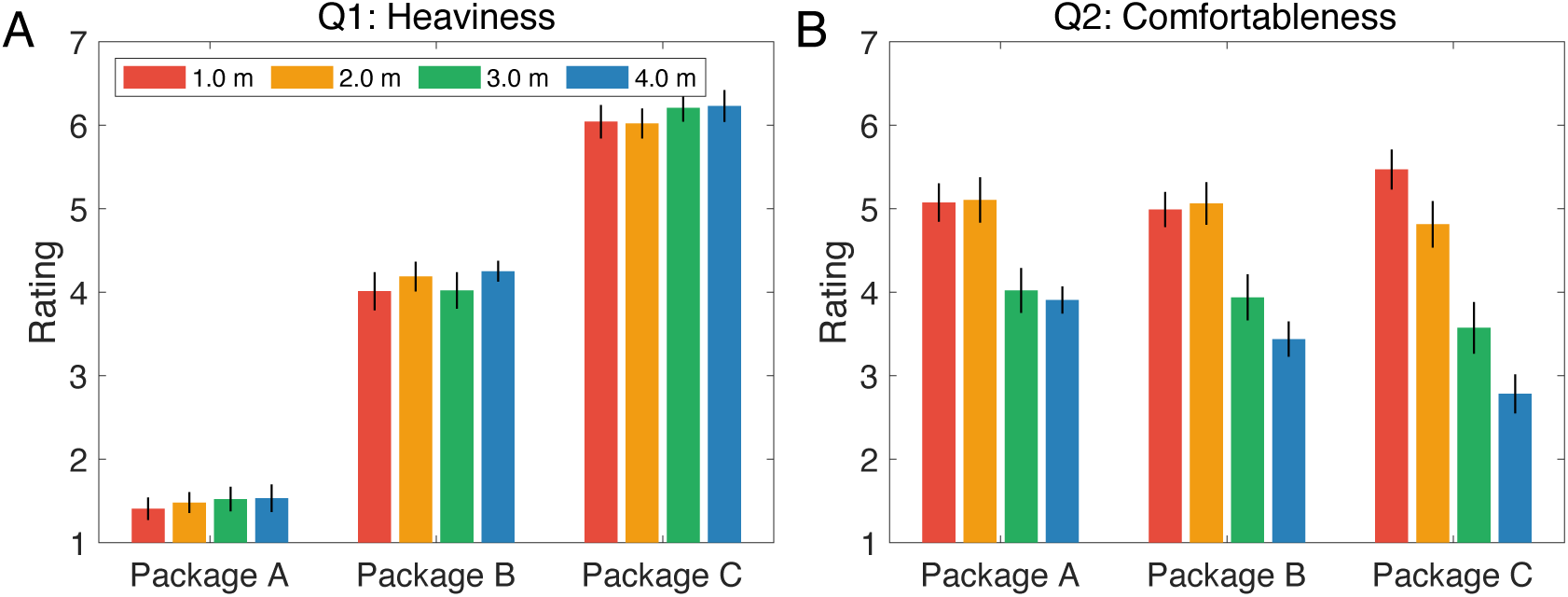
Subjective responses related to **A** package heaviness from 1 (very light) to 7 (very heavy) and **B** degree of comfort with AMR behavior from 1 (very uncomfortable) to 7 (very comfortable). The bars indicate the standard errors.

Fig. 7B shows the average rating score for comfort obtained from the responses to question Q2. Two-way repeated-measures ANOVA showed main effects of distance (*F*(3, 45) = 30.11, *p* < 0.001, partial η^2^ = 0.667) and weight (*F*(1.28, 19.26) = 4.07, *p* < 0.05, partial *η*^2^ = 0.213). In addition, a significant interaction (*F*(6, 90) = 5.15, *p* < 0.001, partial *η*^2^ = 0.256) was observed. Post-hoc tests revealed that stop distances of 1 and 2 m were significantly more comfortable than those of 3 and 4 m (1 m and 2 m > 3 m and 4 m; 1 m vs. 2 m, *pholm* = 0.420; 3 m vs. 4 m, *pholm* = 0.087; others, *pholm* < 0.001). When the AMR did not move (i.e., 4 m stop distance), the participants responded that package C felt more uncomfortable than package A (A > C; A vs. C, *pholm* < 0.001), but this did not occur for other stop distances.

### 3.3 Discussion

The participants’ behaviors in Experiment 2 were similar to those in Experiment 1, but they passed the package slightly later when it was heavier (Fig. 5). This finding suggests that a heavier package was passed more slowly and carefully to prevent falls.

The maximum speed was lower when the AMR stop distance was shorter (Fig. 6A). A shorter distance (1 m) also resulted in a higher hesitation probability (Fig. 6B). In Experiment 1, we observed a similar trend of decreasing maximum walking speed to maintain a distance from the AMR, possibly to avoid feeling uncomfortable with the approaching AMR. However, the participants felt more comfortable at shorter stop distances (Fig. 7B). Thus, hesitation is not related to invading personal space, which becomes shorter if a safe margin can be maintained while walking [63].

The subjective response to heaviness showed that the participants clearly perceived differences in the package weights (Fig. 7A). However, contrary to the hypothesis for Experiment 2, the package weight did not affect the participants’ walking speed or hesitation probability (Figs. 6A and 6B). We used a 10.24 kg load in the heaviest case and a 0.87 m package placement height on the AMR. In a previous study, load weights of 2–12 kg and heights of 0.2–1.7 m were evaluated [7], indicating that loading was lighter in our experiment. Thus, the package weights used in our study might not reach a level that influenced walking. We speculate that humans consider load weight and load carrying separately.

The participants felt uncomfortable when passing heavier packages at a stop distance of 4 m, in which case the AMR did not move from its initial position. A previous study reported that holding a heavier box (10.1 kg) made participants physically more uncomfortable than holding a lighter box (1.1 kg) [64]. Hence, discomfort may be related to placing heavy packages on the AMR.

Woods et al. conducted an experiment in which an AMR approached and passed a snack to participants, who preferred front, front-right, and front-left approaches in a task efficiency assessment [65]. Another study asked participants to perform a home tour task with an AMR, wherein the AMR followed the participant, indicated the locations of furniture and objects, and verified those locations [66]. The participants preferred a distance to the AMR that was within the range of proxemics reported by Hall [39]. These reports and our findings have a common point: participants must approach the AMR to complete the task. By contrast, participants feeling uncomfortable by an AMR invading their personal space was reported when the participants independently completed the task without interacting with the AMR [27, 59]. Thus, a task that requires approaching an AMR reduces discomfort.

## 4 General discussion

We conducted two experiments to clarify human behavior and comfort when placing a package on an AMR while a participant approached the robot. The results of Experiment 1 indicated that the participants’ walking speed decreased when the AMR approached faster and closer. In this case, the hesitation probability of the participant temporarily stopping walking occurred more frequently than in other cases. The results of Experiment 2 indicated that the walking speed also decreased, while the participants felt more comfortable when the AMR approached closely. Interestingly, the participants felt less comfortable with heavier weights when the AMR remained still. Overall, different samples were included in Experiments 1 and 2, and the maximum walking speed (Figs. 3 and 6A) and hesitation probability (Figs. 4B and 6B) exhibited similar trends. Thus, these behaviors seem robust and explainable by biological features.

The participants felt more comfortable when the AMR approached more closely (Fig. 7B). This may be explained as the participant regarding the AMR approach as assistance for load carrying. This is supported by the results indicating that participants’ comfort decreased with heavier packages and longer walking, confirming their desire for the AMR to come closer for assistance. Moreover, although the participants responded with lower comfort for heavier packages when the AMR remained still (i.e., 4 m stop distance), the other stop distances showed similar comfort levels regardless of the package weight. These findings suggest that when participants thought that they had no assistance from the AMR, their comfort decreased. We may apply these findings to correctly control AMRs for loading tasks. For example, an AMR should display a highly cooperative attitude toward a person through a close approach or other types of signals.

Previous studies have reported that personal space (45–80 cm) determined by Hall’s proxemics is preferred when humans and robots communicate with each other or collaborate to complete a task [27, 66, 67]. The range of the corresponding distances was shorter when the participants and AMR were closer (1 m) than the stop distances considered in this study. This suggests that the AMR did not invade the personal space of the participants, and they did not feel discomfort. Nevertheless, this does not directly explain why the AMR proximity decreased the walking speed or increased the hesitation probability in terms of personal space. The distance between humans and robots has also been discussed from the viewpoint of participants’ traits [68] and habituation, as revealed by a study conducted over several weeks [69–72]. In addition, the preferred distance changed depending on the experience with the robots [73]. We conducted the experiments assuming that not all participants were habituated to the AMR because they were naïve to collaborative tasks with AMRs. If we repeatedly ran an experiment over a long period, the perceived heaviness and comfort levels might have changed.

In our experiments, the participant and AMR approached each other (Fig. 1A). In terms of approaching robots, humans prefer the front, front-right, or front-left direction, whereas approaching from the back elicits discomfort [24, 29, 65, 74]. By contrast, even when a robot approaches directly, it can elicit discomfort when the person is sitting [75]. These cases are conceivable. Thus, it is necessary to carefully set up experiments and perform further investigations aimed to reduce discomfort in participants.

If the AMR moved fast, the perceived safety and comfort decreased. A study reported that the acceptable distance for an approaching robot increased when it moved fast [76], suggesting that humans can flexibly adjust safety by maintaining a certain distance. Notably, the estimated distances reported in previous studies were measured as subjectively acceptable for the participant to stop the robot, which differs from the evaluation of the AMR operation.

We can infer the AMR operation in a situation such as that evaluated in this study. AMRs should not cause humans to hesitate when approaching. Our findings indicated that closer proximity increased the hesitation probability and decreased the maximum walking speed. When we regard these changes as harmful to human walking, we identify a tradeoff between these behaviors and the subjective comfort elicited by the AMR. Thus, if we add the AMR speed to a subjective evaluation of AMR operation, we may clarify the perceived safety and comfort.

This study has various limitations. First, we investigated the approaching between a person and AMR considering that they directly faced each other in Experiments 1 and 2. The behaviors may considerably change in unrestricted scenarios, such as when the AMR moves not only forward but also sideways and more freely. We must implement a more flexible motion algorithm in the AMR toward its real-world applicability. For example, a human can pass a package to an AMR that is programmed to stop approaching and avoid humans based on sensor data. However, the experimental paradigm should be carefully designed because the robot’s motion may become unpredictable or uncertain, possibly increasing discomfort [77, 78]. Second, we evaluated only three package weights up to approximately 10 kg. Although the weight did not show a significant effect on the maximum walking speed or hesitation probability (Figs. 6A and 6B), a much heavier package would likely affect the experiment outcome. Third, we did not employ various AMR parameters, such as size and appearance [22, 37, 79], acceleration [76], sound [67], and moving and approaching directions [38, 80]. Rubagotti et al. noted that these factors have been extensively shown to affect perceived safety and comfort [27]. Therefore, further investigations are required to clarify human comfort when interacting with AMRs.

## 5 Conclusion

Although AMRs are expected to flourish in urban environments to improve logistics, it remains unclear how humans behave and pass objects to AMRs when they approach each other. Thus, we investigated how both human behavior and comfort change depending on the stop distance, speed, and weight of a human passing a package to an approaching AMR. When the distance between the participant and AMR was smaller, the walking speed decreased, while the hesitation probability increased. Unexpectedly, the participants felt more comfortable when the AMR approached closer. Additionally, although the weights considered in this study did not affect the participants’ behavior, they felt uncomfortable when the AMR remained completely still. This suggests that the approaching AMR was regarded as assistance in load carrying. To achieve comfortable interactions in load carrying from humans to AMRs, we suggest that the AMR should approach closely as long as humans do not feel personal space invasion.

## Author contributions

Conceptualization: Hideki Tamura; Data curation: Hideki Tamura, Taiki Konno; Formal Analysis: Hideki Tamura, Taiki Konno; Funding acquisition: Hideki Tamura, Shigeki Nakauchi, Tetsuto Minami; Investigation: Hideki Tamura, Taiki Konno; Methodology: Hideki Tamura, Taiki Konno; Project administration: Hideki Tamura, Shigeki Nakauchi, Tetsuto Minami; Resources: Hideki Tamura, Shigeki Nakauchi, Tetsuto Minami; Software: Hideki Tamura, Taiki Konno; Supervision: Hideki Tamura, Shigeki Nakauchi, Tetsuto Minami; Validation: Hideki Tamura, Taiki Konno, Shigeki Nakauchi, Tetsuto Minami; Visualization: Hideki Tamura, Taiki Konno; Writing – original draft: Hideki Tamura, Taiki Konno; Writing – review & editing: Hideki Tamura, Taiki Konno, Shigeki Nakauchi, Tetsuto Minami.

## Ethical Approval

This study was approved by the Institutional Review Board of Toyohashi University of Technology.

## Informed Consent

Informed consent was obtained from all the participants enrolled in this study.

## Data availability

The datasets generated during and/or analyzed during the current study are available from the corresponding author upon reasonable request.

## Acknowledgments

This study was based on the results obtained from project JPNP20004, which was subsidized by the New Energy and Industrial Technology Development Organization (NEDO). This work was supported by the Casio Science Promotion Foundation (39-55).

